# Synergistic effects of predation and parasitism on competition between edible and inedible phytoplankton

**DOI:** 10.1101/2023.05.01.538997

**Authors:** Minoru Kasada, Patch Thongthaisong, Sabine Wollrab, Silke Van den Wyngaert, Christine Kiel, Stella A. Berger, Hans-Peter Grossart

**Affiliations:** Graduate School of Life Sciences, Tohoku University 6-3, Aoba, Sendai 980-8578 Japan; Department of Plankton and Microbial Ecology, Leibniz Institute of Freshwater Ecology and Inland Fisheries, Zur alten Fischerhuette 2, 16775, Stechlin, Germany; Institute of Biochemistry and Biology, Potsdam University, Maulbeerallee 2, 14469 Potsdam, Germany; Berlin-Brandenburg Institute of Advanced Biodiversity Research (BBIB), Germany; Department of Biology, University of Turku, Vesilinnantie 5, 20014 Turku, Finland

**Author notes:** Corresponding author E-mail addresses (MK), (HPG).

**Keywords:** microcosm, aquatic fungi, competition, host-parasite interaction, predator-prey interaction, mycoloop, population dynamics, nutrient effects, community composition, chytrids

## Abstract

Fungi can affect aquatic ecosystems through syntrophic and parasitic interactions with other organisms and organic matter. In pelagic systems, fungal parasites on phytoplankton can control trophic interactions and food-web dynamics, e.g., zooplankton grazing on fungal parasite zoospores creates an alternative energy pathway (termed “mycoloop”) from otherwise inedible phytoplankton species. We aim to investigate how the mycoloop influences community dynamics in aquatic food-webs combining experimental and modelling approaches. We assembled an experimental system consisting of an inedible (host) phytoplankton species and its parasitic chytrid, an edible (non-host) phytoplankton species, and a zooplankton grazer. Chytrids parasitizing increased edible phytoplankton abundance, while zooplankton grazing decreased edible phytoplankton abundance. In the presence of zooplankton and chytrids, competition effects between edible and inedible phytoplankton species depended on nutrient levels. At high nutrient levels, competition was balanced by an indirect positive chytrid effect and negative zooplankton grazing effects on edible phytoplankton. In contrast, at low nutrient levels, we found chytrid had a negative impact on edible phytoplankton synergistically with zooplankton. Mathematical investigations suggest that the synergistic effect can be caused by the mycoloop. This indicates that the mycoloop substantially affects predator-prey interactions and phytoplankton competition with yet unknown ecological consequences.

## Introduction

Fungi including “dark matter” fungi [1] can substantially affect aquatic ecosystems via a range of syntrophic to parasitic interactions with other living organisms and organic matter [2–5]. In pelagic systems, especially zoosporic *Chytridiomycota* (chytrids) represent important phytoplankton parasites which can control food-web dynamics and thus organic matter and energy flow [4] with so far largely unknown ecological consequences. For example, zooplankton grazing on the free-living zoospores of chytrids emerging from infected phytoplankton hosts, which would otherwise be inaccessible for the zooplankton, creates an alternative trophic pathway termed the “mycoloop” [6–8]. Thereby, nutrients within the inedible host cell can be transferred to zooplankton through the fungal zoospores [6,7]. The mycoloop, therefore, increases the efficiency by which energy is transported from lower to higher trophic levels through changes in species interactions and potentially affects community composition [8,9].

Despite their evident importance, knowledge on the role of parasitic fungi in aquatic food-webs is still limited [10].

The mycoloop involves two different types of species interactions, i.e., host-parasite and predator-prey. This is in line with community ecology theory, after which communities with multiple interactions provide the key to understanding community dynamics [11]. Recently, some theoretical expectations regarding the effects of species interactions on community composition have been formulated. For instance, communities with various species interaction types, such as predator-prey, mutualism, or competition, show different community stability features [12], whereby a moderate mixture of antagonistic and mutualistic interactions can stabilize population dynamics [11]. Furthermore, the inclusion of host-parasite interactions can result in differences in population dynamics, including chaotic dynamics in predator-prey systems [13,14]. Results from these theoretical investigations on multiple species interactions show that ecological consequences, e.g., community composition and stability, depend on the respective food-web context and assumed interaction type. Specifically, predictions of community composition and stability may significantly differ from subsystems that only consider one level of specific interaction type (i.e., predator-prey, host-parasite, or competition-mutualism). Thus, an experimental system including the combined effects of different species interaction types should be investigated.

Previous experimental studies in the context of the mycoloop have revealed that zoospores produced by chytrids constitute a good food source for zooplankton, potentially improving their growth [6,15]. In addition, if inedible phytoplankton becomes infected, the produced fungal zoospores can provide an alternative food source for zooplankton as they couple primary and secondary production through the mycoloop [16]. Thus, the mycoloop potentially affects population dynamics and composition of plankton communities not only directly but also indirectly via changing the energy transfer pathway [17]. To our knowledge, no empirical study has directly demonstrated how the mycoloop affects plankton community composition and dynamics in an experimental community. Although theoretical studies have shown that different types of species interactions are key to understanding community stability, there are only a few experimental studies following in detail the population dynamics with more than two types of species interactions [11, 12].

In the present study, we address these knowledge gaps by combining experimental and modeling approaches at different levels of food-web interactions: resource competition, host-parasite, predator-prey, and combinations including the mycoloop. In general, the parasitic chytrid infected the inedible phytoplankton, and their zoospores could be grazed by zooplankton; this trophic link formed the mycoloop. Edible and inedible phytoplankton species competed for a shared nutrient pool, and system dynamics were observed at two different nutrient levels. Thus, depending on the respective nutrient levels, we could experimentally demonstrate how the mycoloop affects phytoplankton competition and how the effect transfers to different trophic levels. Additionally, to reveal the effects of the presence or absence of species interactions on our focal species, we compared the dynamics of the food-web and its individual components (Fig. S1). In this study, four experiments were conducted: (1) simple nutrient competition between the two phytoplankton species (competition-food-web), (2) competition with zooplankton (predation-food-web), (3) competition with chytrids (infection-food-web), and (4) competition with both zooplankton and chytrids (i.e., with the mycoloop, entire food-web). By comparing the community dynamics with competition, host-parasite, and predator-prey interactions in the different food-web subsets, we can provide an ideal example of population dynamics in a community with different, but well-defined types of species interactions. Recent developments in chytrids-phytoplankton cultivation and monitoring methods, now, enable the establishment of experimental community systems [18]. Our experimental community consisted of an inedible phytoplankton species and its parasitic chytrid, an edible phytoplankton species, and a zooplankton species (Fig. S1). As inedible phytoplankton species we used *Staurastrum* sp. and its host-specific parasitic chytrid *Staurastromyces oculus* [18]. *Cryptomonas* sp. served as the edible phytoplankton species and *Daphnia magna* as zooplankton top predator. *Daphnia magna* grazed upon *Cryptomonas* (cell size 5–12 μm), but hardly fed upon *Staurastrum* (cell size 15–45 μm (see Method). Importantly, *Daphnia magna* can directly consume *Cryptomonas* and the chytrid (zoospores) via the mycoloop.

Finally, we developed a mathematical model for our experimental system based on the model of Miki et al. [8] and parameterized the model using data from the experiments, and compared the model predictions with the experimental results. In the experiment, chytrids parasitizing positively affected edible phytoplankton, however, in the presence of zooplankton and chytrids, we found chytrid sometimes had a negative impact on edible phytoplankton synergistically with zooplankton. We investigated how this synergistic effect occurs in the mathematical model systems, with or without the mycoloop for a potential range of parameters and reveal how the mycoloop leads to the results observed in the experiment. Our study indicates that chytrids can substantially affect predator-prey interactions and phytoplankton competition, hinting to potential synergistic effects of predators and parasites on the proliferation of inedible phytoplankton. Therefore, the mycoloop affects the entire community dynamics and these effects differ in dependence on ambient nutrient levels.

## Results

### Community dynamics in the experimental systems

In the simple phytoplankton competition experiment (**competition-food-web**, Fig. S1a), cell density of edible *Cryptomonas*, the preferred zooplankton food, was higher than of inedible *Staurastrum*, which cannot be grazed by zooplankton (Fig. 1a, e). Inedible *Staurastrum* grew better under high-nutrient conditions (Fig. 1e) than under low-nutrient conditions (Fig. 1a), whereas edible *Cryptomonas* showed similar growth patterns independent of the nutrient conditions (Fig. 1a, e). With parasites only (**infection-food-web**, Fig. S1b), cell density of edible *Cryptomonas* was higher than of inedible *Staurastrum*, which was unable to grow due to a high level of lethal chytrid infections (Fig. 1b, f). In this infection-food-web, phytoplankton dynamics were similar under high- and low-nutrient conditions. With predators only (**predation-food-web**; Fig. S1c), the cell density of edible *Cryptomonas* was largely diminished due to zooplankton grazing, whereas cell density of inedible *Staurastrum* was slightly lower than in the simple competition-food-web at low nutrient conditions (Fig. 1c, g). Yet, under high nutrient conditions, the cell density of inedible *Staurastrum* was higher than of edible *Cryptomonas* (Fig. 1g), and this pattern was not visible in any of the other experimental setups. *Daphnia* increased to approximately 100 ind L^-1^, then decreased and became extinct at the end of the experiment (Fig. 1g). Moreover, we observed an additional pattern under high-nutrient conditions, even for the same experimental setup (Fig. 1i). For this pattern, *Daphnia* could not suppress the edible *Cryptomonas* (Fig. 1i). Since we wanted to investigate the combined effect of predation and parasitism, we mainly focused on the “strong impact” (Fig. 1g) pattern, where *Daphnia* was able to suppress the edible *Cryptomonas*.

**Figure 1.**
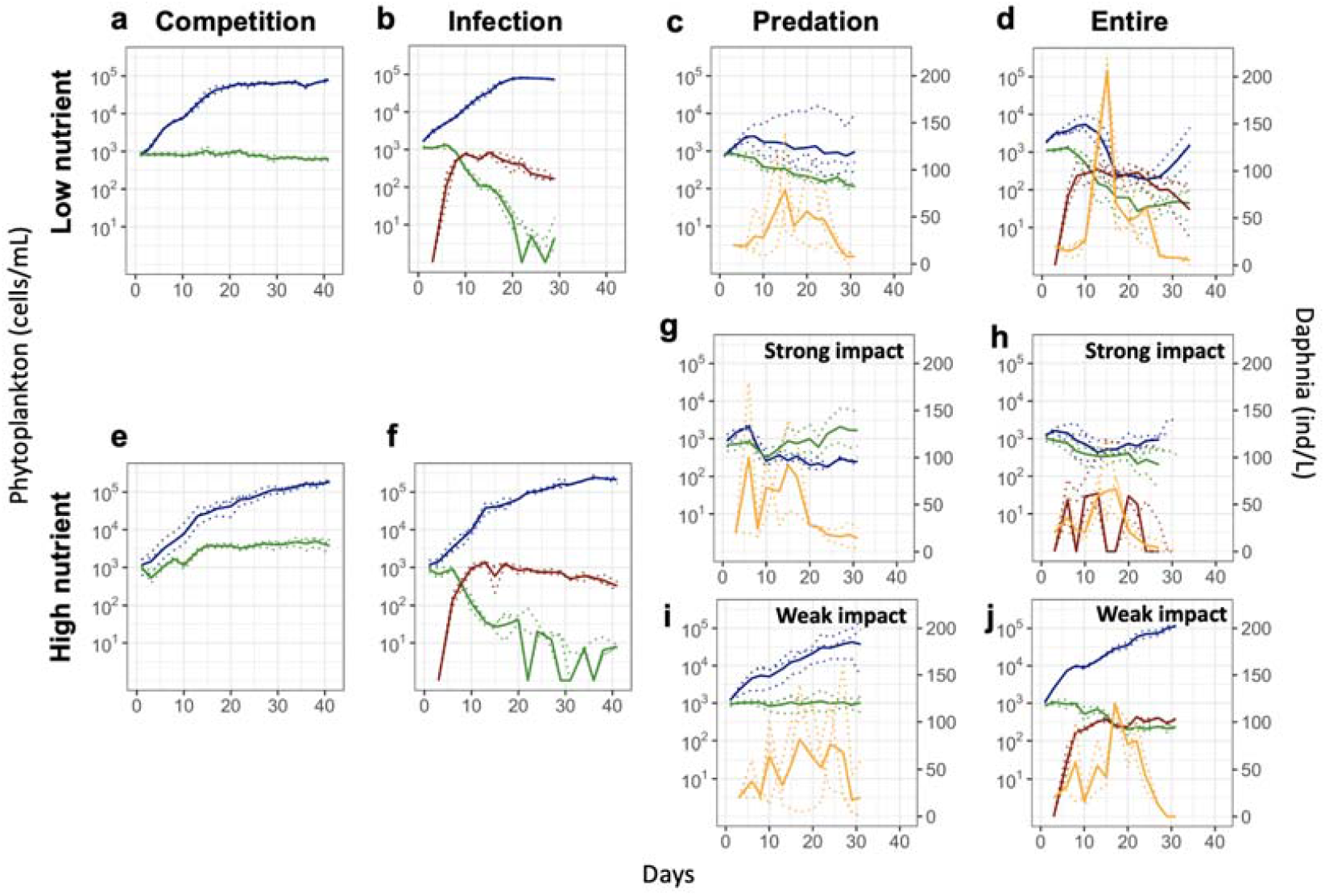
Dynamics of edible *Cryptomonas* and inedible *Staurastrum* in a (a, e) competition system, (b, f) with parasitic chytrids on inedible *Staurastrum*, (c, g, i) with *Daphnia magna*, and (d, h, j) with both parasitic chytrids on *Staurastrum* and *Daphnia magna*. Panels (a-d) represent low nutrient conditions and panels (e-j) represent high nutrient conditions. The experimental setups of (g) and (h)are the same as (i) and (j), respectively. Dotted lines show each replication and continuous lines represent the average of the replications. Blue lines represent the edible phytoplankton, i.e., *Cryptomonas*, green lines represent (inedible) susceptible phytoplankton, i.e., *Staurastrum* in the system without or with chytrids. Red lines represent the number of infected *Staurastrum* cells in the system with chytrids, and orange lines represent the density of *Daphnia*. Note that the y-axis for the abundance of phytoplankton is log scaled.

Finally, with both predators and parasites (**entire food-web**, Fig. S1d), uninfected inedible *Staurastrum* decreased (77.27 cells ml^-1^) and infected *Staurastrum* cells increased (74.24 cells ml^-1^) at the end of the experiment under low nutrient conditions. Edible *Cryptomonas* showed a cubic trend under low nutrient conditions, exhibiting an increase of up to 5,961 cells ml^-1^ (average, day 1–10 in Fig. 1d), followed by a decrease to 193 cells ml^-1^ (average, day 10–24 in Fig. 1d) before cell numbers increased again (day 24–34 in Fig. 1d). The density of *Daphnia* in the entire food-web under low-nutrient conditions reached a very high peak (Fig. 1d), while under high-nutrient conditions, such a variable and high *Daphnia* density was not observed. Moreover, we observed additional patterns for the same experimental setup as the predation food-web under high-nutrient conditions (Fig. 1h and Fig. 1j). In addition to the weak impact on the predation-food-web, *Daphnia* could not suppress edible *Cryptomonas* (Fig. 1j). Yet, average abundance of infected inedible *Staurastrum* in the weak impact case was higher than in the strong impact case. To investigating the combined effect of predation and parasitism, we mainly focused on the “strong impact” (Fig. 1h) pattern. In the strong impact case, infection levels of chytrids were low, and density of *Daphnia* was lower than that under low nutrient conditions for the entire food-web (Fig. d and h).

To investigate the influence of community structure on phytoplankton competition, we visualized the dynamics of the ratio of edible *Cryptomonas* to total cell densities of phytoplankton over time (Fig. 2). Under high nutrient conditions, relative abundance of *Cryptomonas* increased in the infection-food-web, whereas there was a notable decline of *Cryptomonas* for the predation-food-web (Fig. 2a). The system with the entire food-web showed intermediate results between those of the infection- and the predation-food-web (Fig. 2a). We call this the “additive effect” because the entire food-web shows a midway result between the predation-food-web, which is advantageous to the inedible *Staurastrum*, and the infection-food-web, which is advantageous to the non-infective, edible *Cryptomonas*. Under low nutrient conditions, the ratio showed similar patterns to the high nutrient condition for all experimental setups until day 6 (Fig. 2b). Thereafter, edible *Cryptomonas* maintained a high cell number relative to the total phytoplankton number in both the infection- and predation-food-webs; However, the ratio of edible *Cryptomonas* to total phytoplankton in the entire food-web was lower than that in the infection- and predation-food-webs. We call this a “synergistic effect” because the entire food-web does not show the simple midway result between the infection- and predation-food-web. The difference between the ratios was maximized at approximately day 22. At the end of the experiment, the ratio of edible *Cryptomonas* in the entire food-web reached the same value as in the predation-food-web. (Fig. 2b).

**Figure 2.**
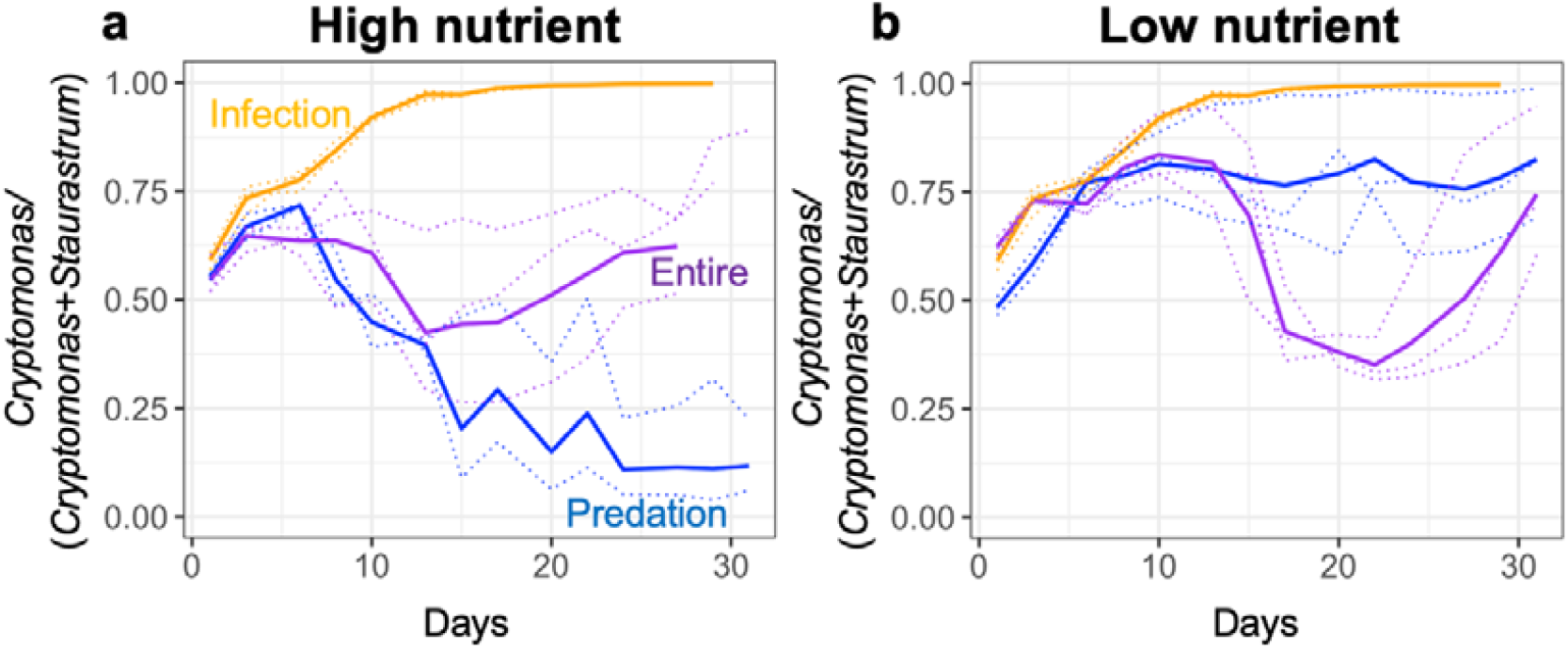
Ratio of edible phytoplankton (*Cryptomonas*) to total phytoplankton cell density over the time course of the experiment under (a) high and (b) low nutrient conditions for the infection-foodweb (orange line), the predation-food-web (blue line), and the entire food-web (purple line). Dotted lines represent replications and solid lines the geometric averages for each model community system.

To explain how this synergistic effect occurs, we analyzed a differential equation model based on Miki et al. [8]. Here, we specifically focused on the phytoplankton competition results (i.e., relative *Cryptomonas* abundance). We first prepared models with all food-web components (i.e., competition-, infection-, predation-, and entire food-web) as we have used in our experimental setups (Fig. S1). Moreover, we tested how the presence of the mycoloop affects the competition results following the transient dynamics of the corresponding mathematical model. For comparison with the experimental results, we focused on model and experimental competition results on day 20 because for both high and low nutrient conditions, differences between the food-web systems were most pronounced between days 15–25 (Fig. 2). To accomplish this, we classified the competition results of systems in the model based on the ratio of edible *Cryptomonas* to the total phytoplankton density, as in the experiment (see Methods).

We prepared two distinct food-web configurations for the entire food-web to investigate the effect of the mycoloop on the competition results. For this, we implemented models with the entire food-web (i) in presence versus (ii) in absence of the mycoloop (no zooplankton ingestion of chytrids). As several physiological parameters remain unknown, we investigated the changes in system dynamics varying specific parameters within biologically reasonable ranges and investigating the results using phase diagrams for all possible pairwise parameter combinations (Fig. 3, Fig. S3-S6). These results confirm that the presence of the mycoloop likely results in a synergistic effect (Fig. 3). This effect is not observed for the entire food-web in the absence of the mycoloop (Fig. 3, Fig. S3-S6). However, occurrence of the synergistic effect also depends on the choice of parameter values. We found some parameter spaces where the synergistic effect occurs under low-nutrient and some where the additive effect occurs under high-nutrient conditions (Fig. 3).

**Figure 3.**
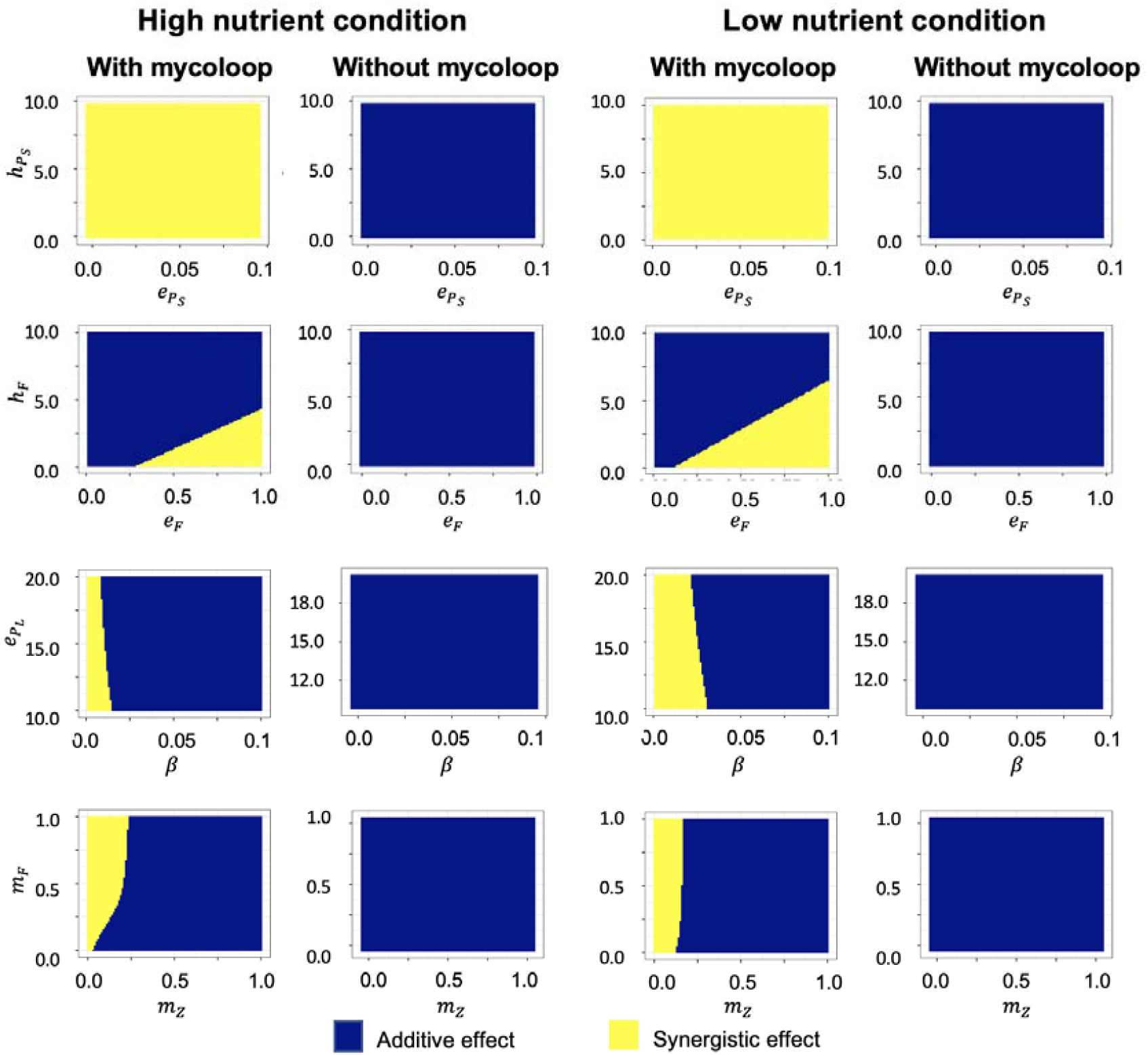
Phase diagrams of the competition results on day 20, indicated by the relative contribution of edible phytoplankton to the total phytoplankton density, for different pairwise parameter combinations under low nutrient conditions (two right columns) versus high nutrient conditions (two left columns) for the entire food-web in presence (first and third column) versus absence (second and fourth column) of the mycoloop. Yellow and blue areas indicate synergistic and additive effects, respectively. The first row represents phase diagrams between conversion efficiency from edible phytoplankton to zooplankton 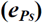 and handling time of zooplankton for edible phytoplankton 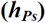. The second row represents phase diagrams between conversion efficiency from chytrids to zooplankton (*e*_*F*_) and handling time of zooplankton for chytrids (*h*_*F*_). The third row represents phase diagrams between infection strength of chytrids (β) and conversion efficiency from inedible phytoplankton to chytrids (*e*_*PL*_). The fourth row represents phase diagrams between zooplankton mortality (*m*_*z*_) and chytrid mortality (*m*_*F*_). The parameter values are given in Table S2.

## Discussion

To understand the single and combined effects of predation and parasitism on phytoplankton resource competition and growth, we compared the population dynamics of different species compositions and combinations of interaction types. We found that the combined effects of predation and parasitism on phytoplankton communities differed depending on the nutrient levels (Fig. 2) and physiological parameters (Fig. 3), e.g. they show that zooplankton and chytrids additively or synergistically influence phytoplankton competition results. The laboratory experiment showed Our model analysis supports that the effects can be explained by the mycoloop. In the performed laboratory experiment, edible *Cryptomonas* had a higher relative concentration within the infection-food-web than in the predation-food-web (Fig. 2), independent of nutrient conditions. This reflects the direct negative impact of *Daphnia* on *Cryptomonas* by predation, while parasites indirectly have a positive impact on *Cryptomonas* by infecting their competitors (i.e., inedible *Staurastrum*). The relative abundance of Cryptomonas was higher in the infection-food-web than in the predation-food-web under both nutrient conditions (Fig2). This suggests that the top-down control of parasites is more important (stronger) than the bottom-up control of nutrients for phytoplankton competition in the experiment [19]. In contrast, nutrients affected competition in the predation-food web; i.e. at high nutrients, Cryptomonas was out-competed by Staurastrum, which became the dominant species (Fig 2a), whereas under low nutrients Cryptomonas remained the dominant species (Fig2b). Thus, both top-down and bottom-up controls seem to drive phytoplankton competition [20, 21]. The degree of the relative increase of edible *Cryptomonas* in the entire food-web depended on ambient nutrient conditions (Fig. 2). Under high-nutrient conditions, the competition of phytoplankton in the entire food-web showed intermediate results between the infection- and predation-food-web (Fig. 2a), suggesting that predation and parasitism have an additive effect on phytoplankton competition. However, under low-nutrient conditions, for the entire food-web, the contribution of edible *Cryptomonas* to the total phytoplankton was even lower compared to that with only chytrids or *Daphnia* (Fig. 2b). Thus, the combined effects of predation and parasitism for this case can be interpreted as synergistic. This is further supported by the model results, which suggest that the mycoloop plays a critical role for the occurrence of synergistic effects (Fig. 3). The model results show that in presence of the mycoloop, both synergistic and additive effects can occur (Fig. 3) in different ranges of the parameters, while in the absence of the mycoloop, no synergistic effect was observed (Fig. S3-S6).

Previous studies indicated the mycoloop affects plankton communities beyond a single host-parasite or predator-prey interaction [6,7], although we have not known the specific phenomenon. In the current study, we found the effect of mycoloop emerging as the synergistic or additive effect in a zooplankton-phytoplankton community and demonstrated details of the population dynamics with more than two types of species interactions. According to the previous model analysis by [8], the mycoloop has both positive and negative effects on edible phytoplankton; positive by reducing energy transfer from edible phytoplankton to zooplankton due to functioning chytrids zoospores as a good resource for zooplankton and negative through the reduction of parasite density and therefore infection risk via zoospore consumption by zooplankton. We found some parameter spaces where the synergistic effect occurs under low-nutrient and some where the additive effect occurs under high-nutrient conditions (Fig. 3). Thus, the model potentially explains the difference between the synergistic and additive effects from the experimental observations by comparing the model results between high- and low-nutrient conditions at a parameter set.

Under high-nutrient conditions, we observed two different dynamics patterns for the predation- and the entire food-web even in the same experimental setup (Fig. 1g, i, h, j). A “strong impact” scenario, where *Daphnia* could suppress edible *Cryptomonas* and a “weak impact” scenario where, *Daphnia* could not suppress edible *Cryptomonas*. These two scenarios were observed more than twice among replications in our experiment, and such phenomenon caused by slight uncontrollable differences among replications in some experiments [22, 23]. A potential explanation for this observation stems from a stochastic process of *Daphnia* demography and differences among the replicated runs of cultures. In the experimental system, the maximum number of *Daphnia* was 60 individuals per flask (250 ml). Stochastic changes in demography may have affected edible phytoplankton biomass. In addition, the extent of individual *D. magna* consumption varies considerably [24]. Another factor is the possibility of deterioration of water quality via phytoplankton overgrowth, as has been observed for phytoplankton mass blooms in nature [25]. When zooplankton fails to control phytoplankton bloom once, the bloom negatively affects zooplankton by the deterioration of water quality, and the top-down control functions less and less. Thus, the initial slight difference in the degree of control will be magnified. In the experiment, a difference in initial grazing impact by the uncontrollable size variation of zooplankton might produce two different scenarios.

In this study, we focused mainly on short-term dynamics of the investigated model system.

Short-term (transient) dynamics are as crucial as the investigation of long-term predictions (steady states), especially for aquatic plankton communities with fast reproduction rates and, therefore, a fast response to daily and seasonal environmental changes, which might keep community dynamics constantly from reaching a stable state [26–29]. For aquatic systems in temperate regions, the initiation of phytoplankton growth is closely related to seasonally changing environmental conditions and starting conditions at the onset of phytoplankton blooms might differ between years and seasons [28]. Based on these results we suggest that the effect of parasites on phytoplankton blooms changes qualitatively depending on nutrient status, i.e. eutrophication level. Our results once more highlight the importance to take both parasite-host and predator-prey interactions into account for predictions on community response to environmental change. If parasites are ignored, a decrease in zooplankton and increasing dominance of inedible phytoplankton would be predicted [30]. however, when parasites are taken into account, temporal decreases in inedible phytoplankton due to parasite infection may occur especially under high nutrient conditions (Fig. 2a, [31,32]). Moreover, our experimental results indicate that under low nutrient conditions, parasites are likely to enhance the dominance of inedible phytoplankton in conjunction with zooplankton (Fig. 2b) via trophic cascading along the mycoloop [6,33,34]. Therefore, the effects of parasitic fungi on phytoplankton blooms can be both positive and negative depending on the overall community structure (i.e., predation, parasite, or entire food-web), eutrophication level, and species-specific differences in physiological parameters (Fig 3). Our experiment was conducted for a relatively short time in comparison to the adaptation time scale (3-5 generations for *Daphnia*), owing to experimental limitations in the possible observation period. For example, parasitic chytrids can adapt to their host phytoplankton [34], and phytoplankton can adapt to zooplankton predators through genetic [35,36] and plastic changes [37] if the observation period is long enough. Thus, the combined effects of parasites and predators on phytoplankton community dynamics and their genetic diversity may become much more complex through eco-evolutionary dynamics [38,39] than indicated by our short-term experiments with a food-web of reduced complexity. This could be addressed in future model studies embracing species diversity and multiple interaction types in a framework that allows for adaptive dynamics to assess long-term trends.

Recent studies have revealed that next to predators, parasites can have a large effect on aquatic community stability and dynamics, and have highlighted the need for data to assess the impact of parasites on entire plankton communities [40]. Here, we found that predators and parasites can synergistically affect the phytoplankton community. This synergistic effect cannot be assessed from the single effects (predation vs. infection) but stems from a complex interaction of direct and indirect effects via a more complex food-web. Our results also revealed that the occurrence of synergistic and additive effects of parasite and predator interactions on phytoplankton community dynamics is influenced by nutrient availability and physiological parameters. This study clearly shows that parasitic fungi, i.e. chytrids, can critically affect aquatic community dynamics via direct and indirect feedback mechanisms with consequences for phytoplankton community composition. This has potential implications for ecosystem functions in aquatic systems such as primary and secondary production. Furthermore, corresponding to previous theoretical work [11], our experimental results indicate that insight from studies on single interaction types might be misleading for assessing effects of multiple interaction types on community dynamics and its ecological consequences. Our study experimentally demonstrated that the effects of parasitic fungi on community composition are different depending on community composition, nutrient availability and physiological parameters, as has been suggested verbally or theoretically [6-8,17], and we specified the additive and synergistic effects. Thus, our study illuminates the ecological role of “dark-matter” fungi on short-term food web dynamics beyond existing knowledge.

## Materials and Methods

### Preparatory experiments: Consumption of Staurastrum by Daphnia magna

To test whether *Daphnia magna* consumes the inedible phytoplankton *Staurastrum sp*., we conducted a preparatory laboratory experiment. We inoculated the inedible algae *Staurastrum* without (control) and with two starved *Daphnia magna* into a 50 ml tube containing modified CHU-10 medium (hereafter mCHU-10). The initial density of *Staurastrum* was adjusted to 1,000 cells ml^-1^. After inoculation, the cultures were incubated overnight in the dark. Each setup consisted of four replications. After overnight treatment, the average density of *Staurastrum* with *Daphnia* was 9,657.5 cells/ml (standard deviation, *SD* = 525.1) and the average density of *Staurastrum* without *Daphnia* was 9,574.9 cells/ml (*SD* = 983.0), an indication that there was no significant difference between the *Daphnia* addition and control (t-test, *t-value =* 0.149, *p =* 0.888). Thus, we could not obtain any evidence that *Daphnia magna* ingested *Staurastrum*, even though *Daphnia* can theoretically feed on particles of *Staurastrum* size (15–45 μm) [41,42].

### Community model system and microcosm experiments

The host-parasite community model system consisted of the host *Staurastrum* sp., (strain STAU1), and the parasite *Staurastromyces oculus* (strain STAU-CHY3), which were established in a previous study [5]. The predator-prey model system consisted of the predator *Daphnia magna* and the prey *Cryptomonas* sp [43]. The medium (mCHU-10) was based on CHU-10 (modified after [44], see Table S1). For the high nutrient conditions, the mCHU-10 medium was diluted twice, and for low nutrient conditions, the mCHU-10 medium was diluted 10 times.

For these experiments, we used a semi-batch culture system (Fig. S7). At the beginning of the experiment, we inoculated 1,000 *Cryptomonas* and *Staurastrum* cells per mL, which had been cultivated in mCHU-10 medium, into 250 ml mCHU-10 medium in a 300 ml wide-mouth Erlenmeyer flask in the all food-web systems (i.e., competition-, predation-, infection-, and entire). After 3 days, we placed 5 *Daphnia* individuals (starved for a minimum of 24 h) in the predation- and the entire food-web systems and 2,500 chytrid-infected *Staurastrum* cells (i.e., final density 10 infected cells ml^-1^) in the infection- and the entire food-web systems. During the experiment (29-43 days), every two or three days (3 days/week), 50 ml samples were taken from the cultures and 50 ml of fresh medium was added (Fig. S7). When we observed three *Daphnia* or more in the sample (50 ml) we counted them on a stereo microscope and multiplied their number by five to extrapolate our counts to total *Daphnia* density in the experimental flask. Alternatively, we visually counted the entire number of *Daphnia* in the experimental flask from outside the flask. When less than three *Daphnia* were observed in a sample, we returned the sampled *Daphnia* to the flask to avoid the extinction of *Daphnia* by a stochastic process of demography. To monitor phytoplankton cell density, we used image-based flow cytometry (FlowCam CYANO, 8000 series, Yokogawa Fluid Imaging Technologies, Inc., Scarborough, Maine, USA) equipped with a 300 μm flow-cell and a 4 × objective. Each run captured more than 1,000 pictures of phytoplankton species per sample in a flow-through system. Each sample was run twice, and the average value was used. The edible and inedible phytoplankton species, *Cryptomonas* and *Staurastrum*, were automatically classified during the runs by size settings related to area-based diameter (ABD), with 5–12 μm diameter for *Cryptomonas*, and 15–45 μm diameter for *Staurastrum*, and color (*Staurastrum* pictures include blue color). To monitor chytrid infections, we observed a minimum of 100 cells per sample under an inverted fluorescence microscope (Nikon ECLIPSE Ti2, Nikon, Japan) after staining chytrid sporangia with calcofluor-white [45].

Each experimental setup was replicated at least thrice. However, we obtained different dynamics from the same setup for the system with *Daphnia* under high-nutrient conditions (see Results). In these cases, additional experimental rounds were performed to obtain at least two replicates with similar population dynamics. During the experiment, the cultures were maintained under constant conditions in a temperature-controlled room at 19 °C under continuous light (200 lx).

### Model formulation and parameterization

To compare the experimental data with theoretical expectations, we constructed a differential equation model building up on an existing model for the investigated study system [8]

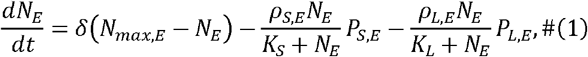

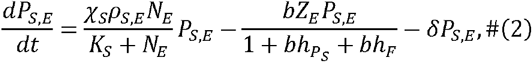

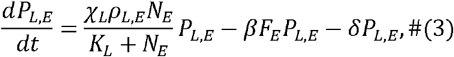

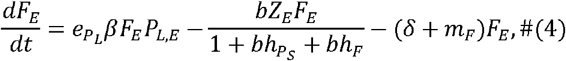

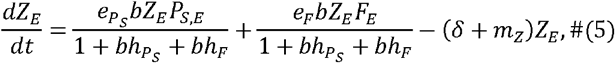

The food-web model reflected the study system and consisted of five state variables (Eqs. 1-5); edible non-host phytoplankton *P*_*S,E*_, (small) and non-edible host phytoplankton, *P*_*L,E*_,(large) competed for a shared resource, *N*_*E*_ assumed to be nitrogen [μgP L^-1^], the parasitic chytrids (a fungal parasite), and zooplankton *Z*_*E*_. The subscript *E* indicates the environmental conditions with high (*E* = *H*) or low (*E* = *L*) nutrient levels. Unlike Miki et al. [8], who measured the state variables (population densities) in units of the limiting nutrient, we measured the state variables in units of cell/individual density to directly compare them with the experimental data. Instead of the experimental semi-batch culture system, nutrient dynamics (Eq. 1) in the model were assumed to follow chemostat dynamics with continuous dilution rate δ = 0.086 day^-1^ reflecting the average exchange rate of the corresponding experiment where 1/5 of the medium was changed every 3 days per week (Eq. 1). To mimick the nutrient concentrations of the experimental set-up, the maximum nutrient availability *N* _*max*_ [μg P L^-1^] was assumed to be 890 μgP L^-1^ and 178 μgP L^-1^ for high and low nutrient cases, respectively. Both phytoplankton species gain energy via nutrient uptake of the limiting nutrient, following Monod kinetics with maximum nutrient uptake rate *ρ*_*S,E*_ carrying capacity *K*_S_ and conversion efficiency *X*_s_ (Eqs. 2,3). Furthermore, phytoplankton experience losses due to the dilution rate d and grazing by *Z*_*E*_ for, *P*_*SE*_ as well as infection by *F*_*E*_ for *P*_*L,E*_ (Eqs. 2,3). The population dynamics of parasitic chytrids (Eq. 4) are determined by infection rate βby assuming linear dependence of infection on host density *P*_*L,E*_. Infected host cells are transferred to chytrid density based on assimilation efficiency 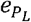. Chytrids experience losses due to dilution rate *d* and background mortality *m*_*F*_ Zooplankton (Eq. 5) gains energy through consumption of, *P*_*S,E*_ and *F*_*E*_ with ingestion following a saturating Holling type-II functional response with maximum ingestion rate *b* and handling time 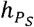 and *h*_*F*_ and assimilation efficiency 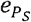 and *e*_*F*_ for small phytoplankton and parasitic chytrids, respectively. Zooplankton experiences losses according to the dilution rate d and a background mortality rate *m*_*z*_.

We used Eqs. 1-5 to investigate the community dynamics of (1) the entire food-web (Eqs. 1-5) as well as for the food-web in the absence of the following: (2) zooplankton (infection-food-web *F*_*E*_) (3) chytrids (predation-food-web *Z*_*E*_), and (4) both *F*_*E*_ and *Z*_*E*_ (competition-food-web). For these sub-foodwebs, the equation system was reduced accordingly and the respective state variables *Z*_*E*_ / *F*_*E*_ was set to zero. To assess if the “synergistic effect” observed in the experiment is caused by the influence of the mycoloop (the direct feeding link from chytrids to zooplankton) on the overall community dynamics, we compared the model results for the entire food-web in presence versus absence of a direct feeding link between chytrids and zooplankton by including the ingestion term of zooplankton for chytrids 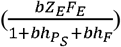 or not, respectively. We classified the competition results of the model systems based on the ratio of edible *Cryptomonas* to the total phytoplankton density, as in the experiment. Notably, *P*_*s, P*_ (*t*) and *P*_*L, P*_ (*t*) were represented as the number of edible and inedible phytoplankton in the predator-prey model, respectively, where *t* is the time (day). Competition between phytoplankton species was measured using the relative contribution of edible phytoplankton to the total prey density *P*_*s, P*_ (*t*) / (*P*_*s, P*_ (*t*) + *P*_*L, P*_ (*t*)) in (i) the predation-food-web *C*_*P*_ (*t*), (ii) the infection-food-web *C*_*I*_ (*t*), and (iii) the entire food-web *C*_*E*_ (*t*). When the “additive effect” happens, the competition results for the different subsystems on day 20 met the following criteria:

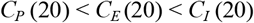

This qualitatively corresponds to the competition results under high-nutrient conditions for the experiment (Fig. 2a). When the “synergistic effect” happens, the competition results for the different subsystems on day 20 met the following criteria:

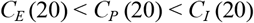

This qualitatively represents the “synergistic effect” and corresponds to the competition results of the low-nutrient experiment (Fig. 2b).

The parameter values for the nutrient uptake of the two phytoplankton species (*ρ*_*S,H*,_ *ρ*_*S,L*_, *ρ*_*L,H*_, *ρ*_*L,L*_,*X*_*S*_,*X*_*L*_ were estimated based on a model fit to the experimental data of the competition experiment (both phytoplankton species in the absence of zooplankton and chytrids). To accomplish this, we investigated the dynamics of the competition-food-web (eq[1-3]). We compared the cell numbers of *Cryptomonas* and *Staurastrum* obtained from the experiment with the density of, *P*_*S,E*_ and, *P*_*L,E*_ that was predicted by the model at the respective sampling (time) points, using log-transformed values. Model fitting was performed using the Levenberg-Marquerdt method available in the “minpack.lm” R package (ver 1.2.1, [46]). We varied the respective parameters to minimize the sum square of the differences between model prediction and observation. The parameter values were estimated at the same time for high or low nutrient condition. Starting conditions for the fitting model were chosen to reflect the average values of the respective state variables on day 1 of the competition experiment. According to the fitting method, (*ρ*_*S,H*,_ *ρ*_*S,L*_, *ρ*_*L,H*_, *ρ*_*L,L*_,*X*_*S*_ *and X*_*L*_ were estimated to be 0.331 μg day^-1^, 0.421 μg day^-1^, 27.1, 19.0 μg^-1^ day^-1^, 0.814×10^6^ cells μg^-1^, and 0.0057×10^6^ cells μg^-1^, respectively (see also Table S2, Fig. S8). All other model parameters were estimated based on available literature (Table S2)

For the simulations, the initial conditions were chosen to reflect the average values of the initial conditions on day 3 of the experiment, when *Daphnia* and chytrids were inoculated and correspondingly the food-web was fully assembled. As only infected cells of *Staurastrum* were added and the exact number of zoospores was not known, the initial density of chytrids was set to 0.15 ind L^-1^, assuming that 0.01 of added *Staurastrum* cells L^-1^ would be converted to 15 zoospores. System dynamics were calculated using the package “deSolve” (ver 1.27.1, [47]) in R (version 4.2.1) (R Core team 2022). The step size of the numerical calculations along time were set to 0.01. Between days 15–25, strong differences between the experimental results were observed under high and low nutrient conditions (Fig. 2); hence, the results simulated for day 20 were used to assess the occurrence of synergistic versus additive effects. This was done by calculating the relative contribution of edible phytoplankton to the total prey density *P*_*s, P*_ (*t*) / (*P*_*s, P*_ (*t*) + *P*_*L, P*_ (*t*)) between the entire food-web in presence versus absence of the feeding link from chytrids to zooplankton. We also checked the results on day 18 and 22, and the conclusion is not affected by the choice of dates.

Furthermore, a sensitivity analysis was performed for all but the directly fitted parameters, varying them within biologically plausible ranges (see table S2) to investigate the model sensitivity to the choice of parameter values and how it affects predictions on the occurrence of synergistic versus additive effects of the mycoloop. For the sensitivity analysis all possible pairwise comparisons between the chosen parameters were performed 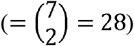 The results are illustrated in phase diagrams (Fig. 3, Fig S3-S6).

## Acknowledgement

This study was supported by a Grant-in-Aid for JSPS Fellows (MK 19J00864) and the German Research Foundation (DFG) Priority Program 1704: DynaTrait (WO 2273/1-1) and (GR 1540/30-1), and funding for instrumentation by the Federal Ministry of Education and Research (BMBF), (funding reference number 033W034A). We thank Dr. Takeshi Miki for his helpful comments, and Mr. Nicholas O’Connor and Editage (www.editage.com) for English language editing.

## Notes

### Competing Interest Statement

The authors have declared no competing interest.

