## Supplementary Information for "Synergistic effects of predation and parasitism on competition between edible and inedible phytoplankton"

**Table S1. Recipe of modified CHU10 (mCHU10) medium.**

Stock solution	mL/Liter	mg/Liter

Na_2_SiO_3_(5H_2_O)	10.824g/250ml	2.5ml/L	43.296mg/L

Ca(NO_3_)_2_(4H_2_O)	2.878g/50ml	1ml/L	57.56mg/L

K_2_HPO_4_(3H_2_O)	0.6551g/50ml	1ml/L	13.1mg/L

MgSO_4_(7H_2_O)	1.25g/50ml	1ml/L	25mg/L

Na_2_CO_3_	1g/50ml	1ml/L	20mg/L

Fe-EDTA	1.85g/50ml	1ml/L	37mg/L

VitaminB12	1mg/1ml,	50μl/50ml	1ml/L	1μg/L

Biotin	1mg/1ml,	50μl/50ml	1ml/L	1μg/L

Thiamine HCl	10mg/50ml	1ml/L	0.2mg/L

pH of the final medium was adjusted to 6.4 for diatoms and green algae.

**Table S2. Parameter, initial values, and descriptions in the model.**

| Parameter | Description | Value | Reference/Range |
| --- | --- | --- | --- |
| $N_{max,H}$ | Concentration of limiting nutrient in the high nutrient medium | 890 μg P/L | Set |
| $N_{max,L}$ | Concentration of limiting nutrient in the low nutrient medium | 178 μg P/L | Set |
| $\delta$ | Dilution rate of the semi-batch system | 0.086 /day | Set |
| $\rho_{S,H}$ | Cryptomonas consumption rate in the high nutrient condition | 0.331 μg/day | Fitted |
| $\rho_{S,L}$ | Cryptomonas consumption rate in the low nutrient condition | 0.424 μg/day | Fitted |
| $\rho_{L,H}$ | Staurastrum consumption rate in the high nutrient condition | 27.1 μg/day | Fitted |
| $\rho_{L,L}$ | Staurastrum consumption rate in the low nutrient condition | 19.2 μg/day | Fitted |
| $\chi_{S}$ | Cryptomonas (10^6^ cells) conversion efficiency | 0.814 10^6^ cells/μg | Fitted |
| $\chi_{L}$ | Staurastrum (10^6^ cells) conversion efficiency | 0.0057 10^6^ cells/μg | Fitted |
| $K_{S}$ | Minimum half-saturation of Cryptomonas | 0.929 μg P/L | (1)* |
| $K_{L}$ | Minimum half-saturation of Staurastrum | 0.929 μg P/L | (1)* |
| $b$ | Daphnia maximum consumption rate | 0.036 L/ind day | (1)* |
| $h_{P_{S}}$ | Handling time of Daphnia for Cryptomonas | 3.09 days | Variable (0.1-10) |
| $h_{F}$ | Handling time of Daphnia for Chytrids | 0.61 days | Variable (0.1-10) |
| $\beta$ | Infectivity strength | 0.314/day | Variable (0.01-0.5) |
| $e_{P_{L}}$ | Conversion efficiency from Staurastrum to the chytrids | 15 spores/cell | (2)* |
| $e_{P_{S}}$ | Conversion efficiency from Cryptomonas to Daphnia | 0.0924 ind/10^6^ cells | Variable (0.001-0.1) |
| $e_{F}$ | Conversion efficiency from Chytrids to *Daphnia* | 0.48 ind/10^6^ spores | Variable (0.01-1.0) |
| $m_{F}$ | Chytrids mortality rate | 0.04 /day | Variable (0-1) |
| $m_{Z}$ | *Daphnia* mortality rate | 0.03 /day | Variable (0-1) |
| $N_{0}$ | Initial nutrient concentration | $N_{max,E}$ μg N/L | Set |
| $P_{S,H}(3)$ | Initial density (day 3) of Cryptomonas in the high nutrient condition | 0.778 10^6^ cells/L | Set |
| $P_{L.H}(3)$ | Initial density (day 3) of Staurastrum in the high nutrient condition | 1.403 10^6^ cells/L | Set |
| $P_{S,L}(3)$ | Initial density (day 3) of Cryptomonas in the low nutrient condition | 0.740 10^6^ cells/L | Set |
| $P_{L,L}(3)$ | Initial density (day 3) of Staurastrum in the low nutrient condition | 0.718 10^6^ cells/L | Set |
| $F(3)$ | Initial density (day 3) of Chytrids zoospore | 0.15 10^6^ spores/L | (2)* |
| $Z(3)$ | Initial density (day 3) of *Daphnia* | 20 ind/L | Set |

* Value is assumed from (1) Miki et al. (2011) and (2) Van den Wyngaert et al. (2017).

**SI text.**

**Calculating phytoplankton volume.**

For *staurastrum,* we approximated the volume using two delta cones. It was assumed that the bottom shape was an equilateral triangle. The volume was calculated as follows:

$$\frac{1}{2}a_{s}\times\frac{a_{s}\sqrt{3}}{2}\times\frac{1}{3}\times b_{s}=\frac{{a_{s}}^{2}b_{s}\sqrt{3}}{12}$$

where *a_s_* is the length of the side of the bottom triangle and *b_s_* is the total height of the two delta cones (Fig S2b). Length data were obtained using the average values of 10 *Staurastrum* individuals. Here *a_s_* and is *b_s_* were 30.18 μm (SD 2.78) and 23.65 μm (SD 1.71), respectively. The volume of *the staurastrum* was calculated to be 3,140.08 μm^3^ (SD 709.53). For *Cryptomonas*, the volume was approximated using an ellipsoid. Thus, the volume was calculated as follows:

$$4\times\pi\times\frac{a_{c}}{2}\times\frac{a_{c}}{2}\times\frac{b_{c}}{2}\times\frac{1}{3}=\frac{{\pi a_{c}}^{2}b_{c}}{6}$$

where *a_c_* is the shortest axis of the ellipsoid and *b_c_* is the longest axis of ellipsoid (see Fig S2d). Length data were obtained using the average values of 10 *Cryptomonas* individuals. Here *a_c_* and is *b_c_* were 5.80 μm (SD 1.13) and 11.49 μm (SD 1.33), respectively. The volume of *Cryptomonas* was calculated to be 208.80 μm^3^ (SD 84.41). Thus, the volume of *Staurastrum* was 15.04 (=3140.08/208.80) times higher than that of *Cryptomonas.*

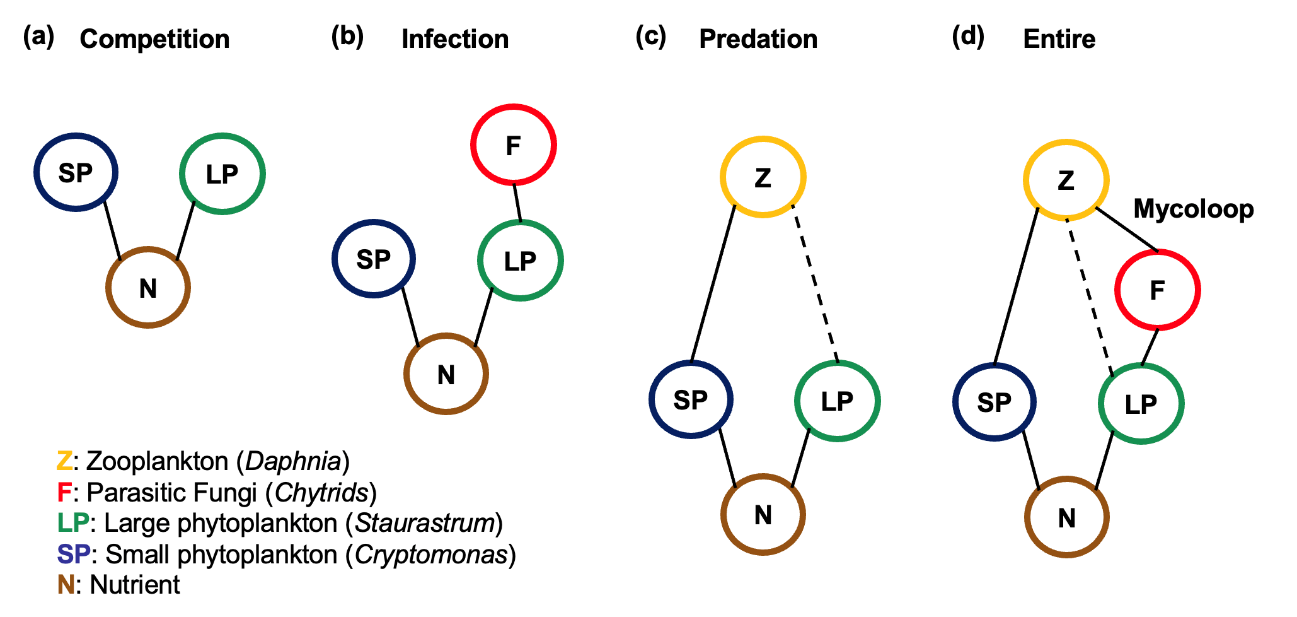

**Figure S1. Model communities used in our experiment systems with (a) resource competition (competition food web), (b) competition and host-parasite interactions (infection food web), (c) competition and predator-prey interactions (predation food web), (d) competition, host-parasite, and predator-prey interactions (entire food web). Brown, blue, green, red, and yellow circles represent nutrient, small phytoplankton, large phytoplankton, fungi, and zooplankton in the system, respectively. Black lines indicate trophic interaction. Dashed lines indicate possible, but empirically unsubstantiated, zooplankton grazing on inedible phytoplankton (see method).**

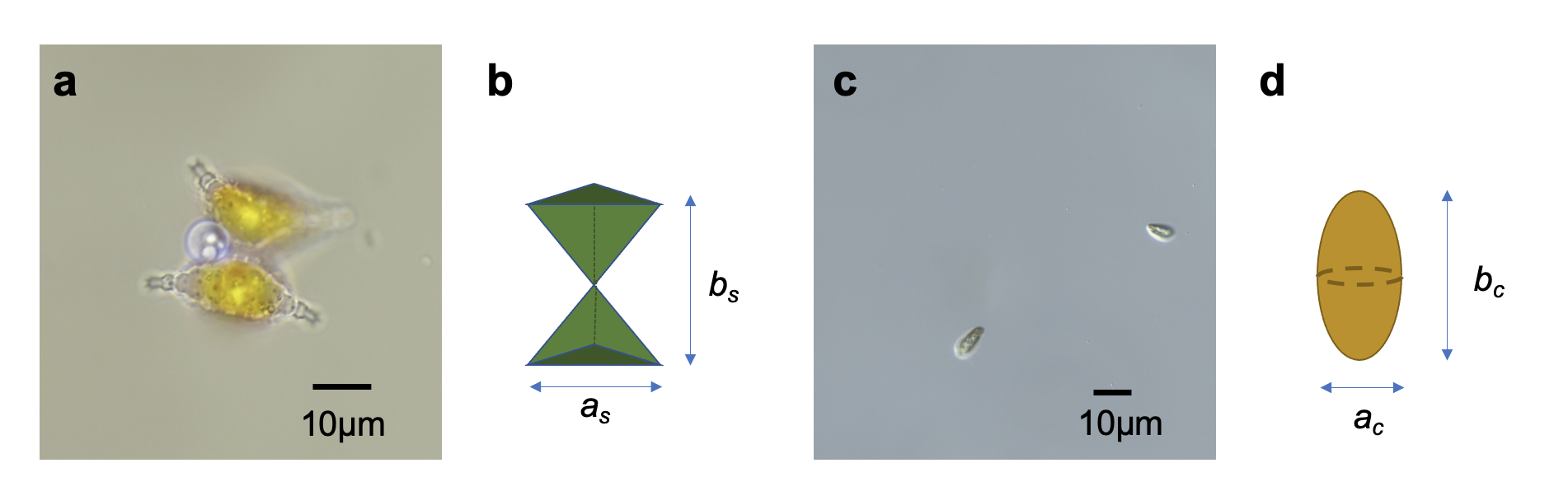

**Figure S2. Images of phytoplankton and measured lengths to calculate their volume.**

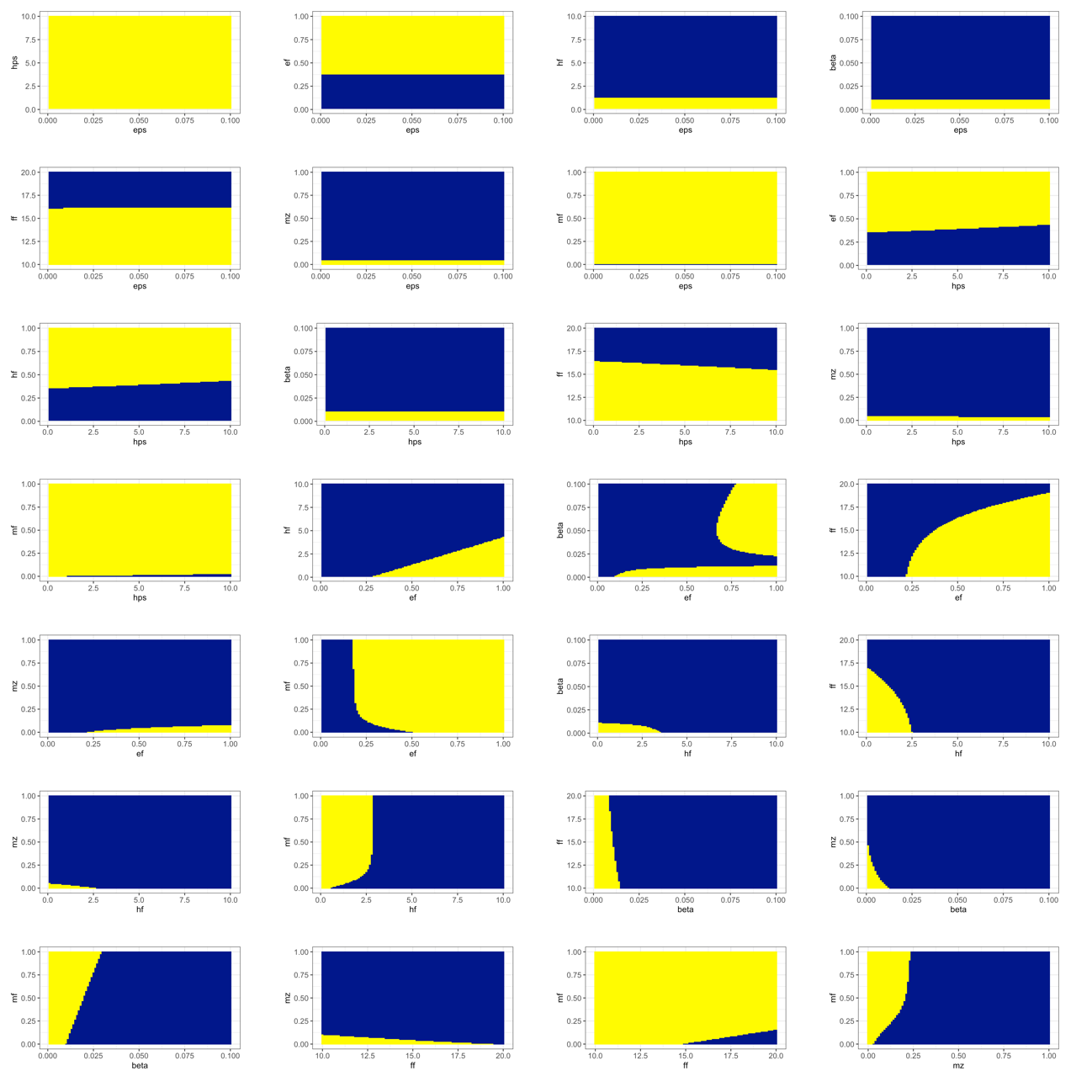

**Figure S3. Phase diagrams of competition results on day 20 for all combination of the parameters in the model with the mycoloop under the high nutrient condition. The other descriptions and parameter values, except for the focal values, were the same as those of Figure 3.**

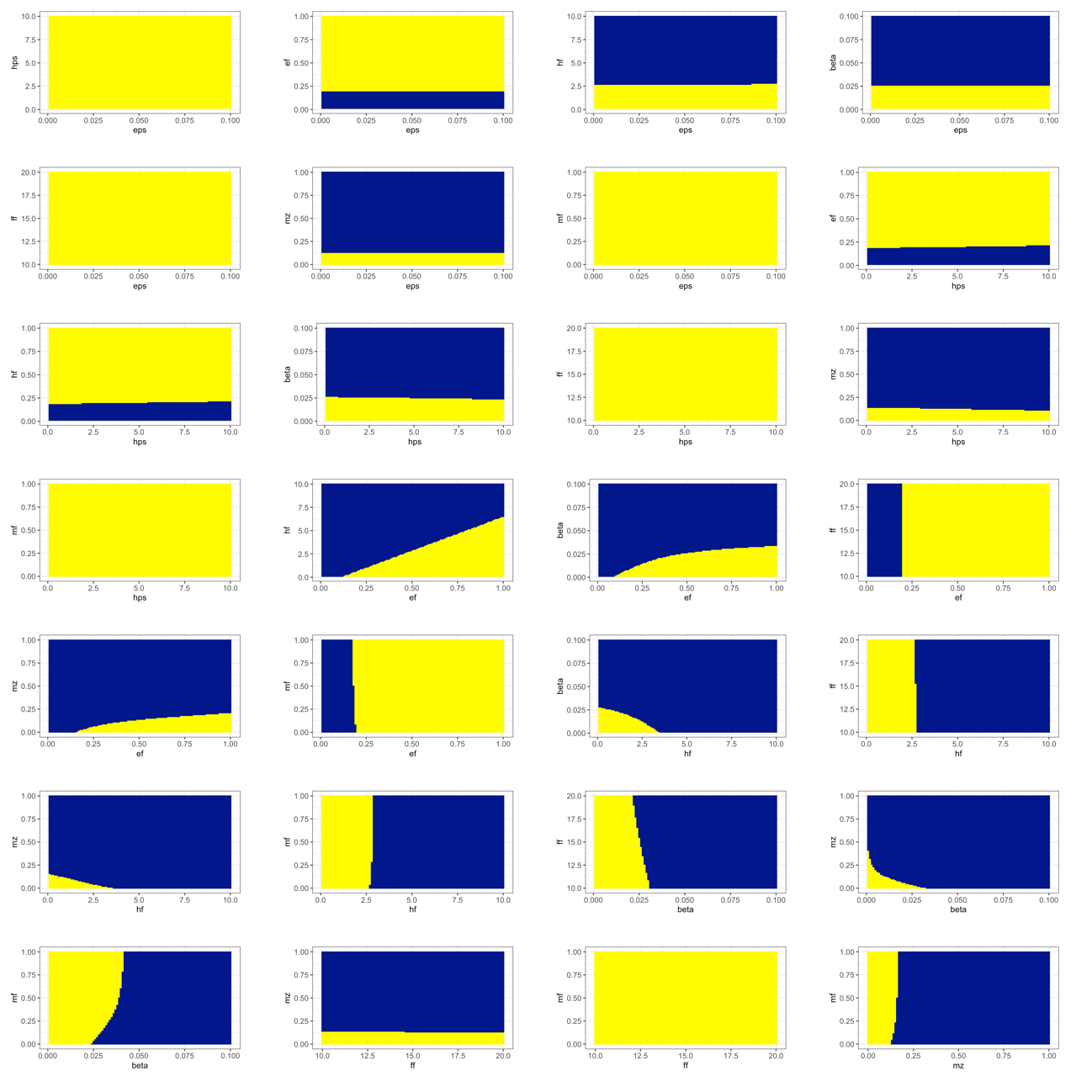

**Figure S4. Phase diagrams of competition results on day 20 for all combination of the parameters in the model with the mycoloop under the low nutrient condition. The other descriptions and parameter values, except for the focal values, were the same as Figure 3.**

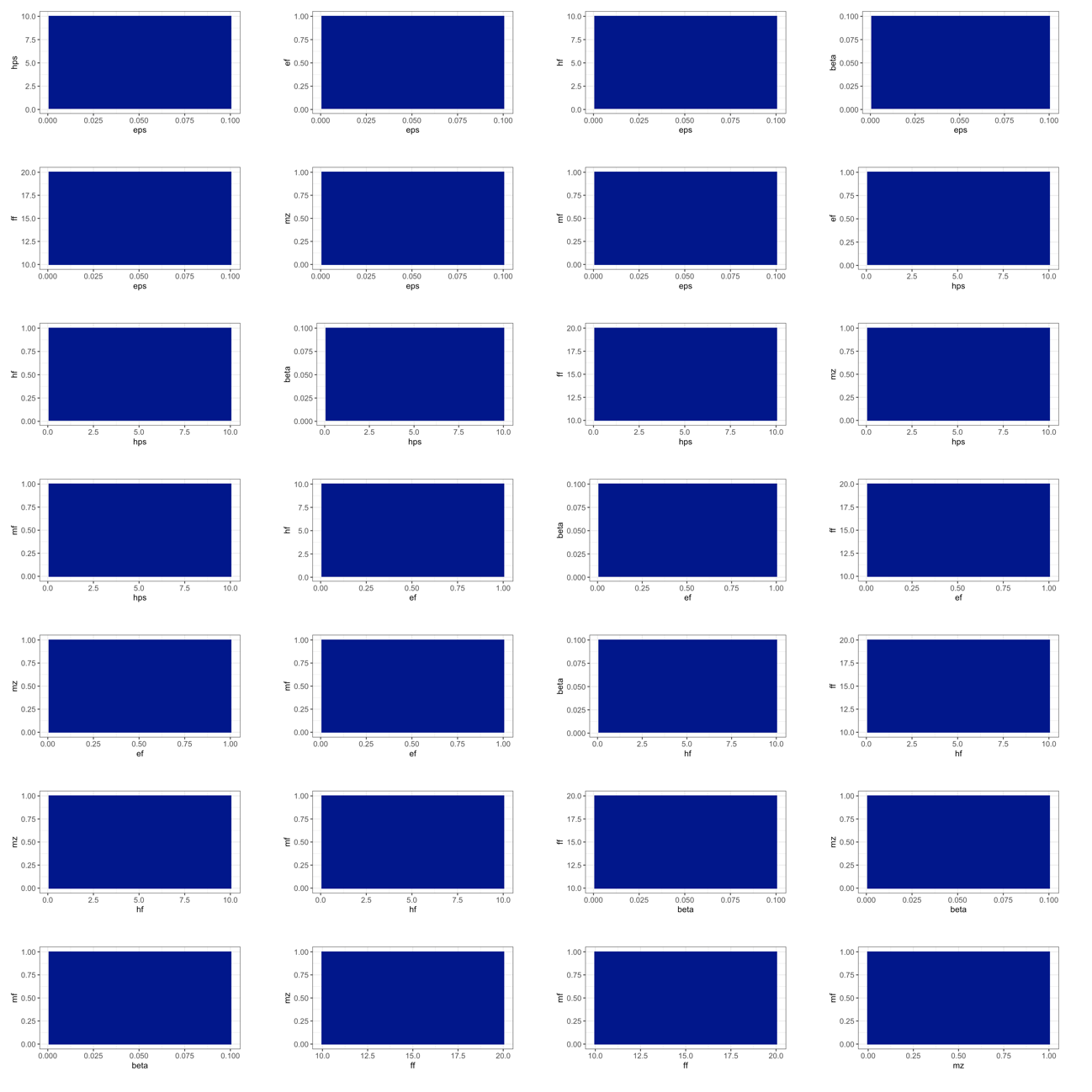

**Figure S5. Phase diagrams of competition results on day 20 for all combination of the parameters in the model without the mycoloop under the high nutrient condition. The other descriptions and parameter values, except for the focal values, were the same as Figure 3.**

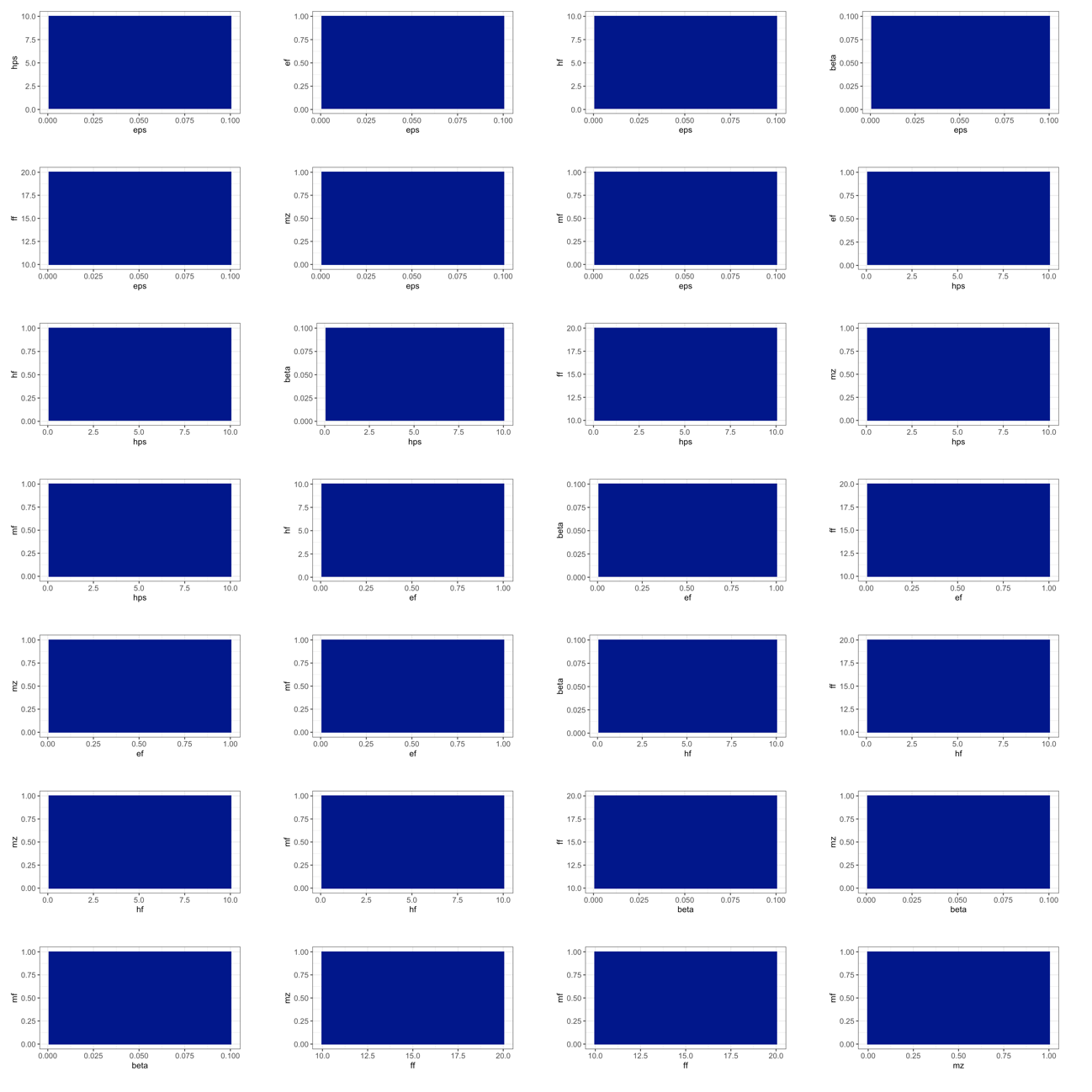

**Figure S6. Phase diagrams of competition results on day 20 for all combination of the parameters in the model without the mycoloop under the low nutrient condition. The other descriptions and parameter values, except for the focal values, were the same as Figure 3.**
